# Real-time control of a hearing instrument with EEG-based attention decoding

**DOI:** 10.1101/2024.03.01.582668

**Authors:** Jens Hjortkjær, Daniel D.E. Wong, Alessandro Catania, Jonatan Märcher-Rørsted, Enea Ceolini, Søren A. Fuglsang, Ilya Kiselev, Giovanni Di Liberto, Shih-Chii Liu, Torsten Dau, Malcolm Slaney, Alain de Cheveigné

**Affiliations:** Hearing Systems Section, Department of Health Technology, Technical University of Denmark, Kgs. Lyngby, Denmark; Danish Research Centre for Magnetic Resonance, Centre for Functional and Diagnostic Imaging and Research, Copenhagen University Hospital - Amager and Hvidovre, Copenhagen, Denmark; Laboratoire des Systèmes Perceptifs, Paris, CNRS UMR 8248, France; Département d’Etudes Cognitives, Ecole Normale Supérieure, Paris, PSL, France; Institute of Neuroinformatics, University of Zurich and ETH Zurich, Zurich, Switzerland; School of Computer Science and Statistics, Institute of Neuroscience, Trinity College, The University of Dublin, Dublin, Ireland; Center for Computer Research in Music and Acoustics (CCRMA), Stanford University, Stanford, CA, United States of America

## Abstract

Enhancing speech perception in everyday noisy acoustic environments remains an outstanding challenge for hearing aids. Speech separation technology is improving rapidly, but hearing devices cannot fully exploit this advance without knowing which sound sources the user wants to hear. Even with high-quality source separation, the hearing aid must know which speech streams to enhance and which to suppress. Advances in EEG-based decoding of auditory attention raise the potential of neurosteering, in which a hearing instrument selectively enhances the sound sources that a hearing-impaired listener is focusing their attention on. Here, we present and discuss a real-time brain-computer interface (BCI) system that combines a stimulus-response model based on canonical correlation analysis (CCA) for real-time EEG attention decoding, coupled with a multi-microphone hardware platform enabling low-latency real-time speech separation through spatial beamforming. We provide an overview of the system and its various components, discuss prospects and limitations of the technology, and illustrate its application with case studies of listeners steering acoustic feedback of competing speech streams via real-time attention decoding. A software implementation code of the system is publicly available for further research and explorations.

## 1. INTRODUCTION

Listeners with normal hearing can follow speech in ‘cocktail party’ scenarios with competing speech sources by selectively attending to relevant speakers (Cherry, 1953; Yost, 1997). Hearing loss greatly reduces this ability, even when the sound input to the ear is amplified by a hearing aid (Bronkhorst, 2000). Consequently, difficulties with communication in everyday noisy situations continue to be the most common complaint from hearing-aid users (Kochkin, 2010; Lesica, 2018). Acoustic amplification or noise suppression in hearing aids can restore some level of audibility, and sound separation algorithms can suppress unwanted sounds to some degree, but the hearing aid cannot help a user follow an individual speech stream among many without knowing which stream the user wants to listen to. Even if the hearing instrument could perform high-quality acoustic scene analysis and separate all the individual sound sources in a complex acoustic environment, it still needs to know which sound sources to enhance and which to suppress. This requires a user input to the hearing aid.

Consequently, the idea of a *cognitively controlled hearing aid* has been proposed to overcome this basic challenge (cocoha.org, Dau et al., 2018; Slaney et al., 2020; Geirnaert et al., 2021). If a user’s selective auditory attention can be measured from brain signals, then the brain-decoded attention signal can be used to control a speech separation process within the hearing instrument. The attention signal would then be used to control the relative levels of competing speakers in real time. Such a hearing instrument would constitute a form of neural prosthesis that assists the hearing-impaired listener by automatically enhancing relevant sound sources in a complex acoustic scene.

The vision of this neuro-steered hearing technology has been fueled by successes in auditory attention decoding from EEG or other neurophysiological signals (O’Sullivan et al., 2015; Mirkovic et al., 2015; Van Eyndhoven et al., 2016; Aroudi et al., 2016; Akram et al., 2016; Miran et al., 2018; Ciccarelli et al., 2019). In offline attention decoding studies, a main direction has been to examine the correlation between continuous EEG responses to speech mixtures and acoustic stimulus features of competing speech streams (Ding and Simon, 2012a; O’Sullivan et al., 2015). The main observation is that low-frequency cortical activity (*<*10 Hz) synchronizes with slow fluctuations in the speech envelope (*<*10 Hz) and that this synchronization is stronger for attended speech streams compared to ignored ones (Mesgarani and Chang, 2012; Power et al., 2012; Ding and Simon, 2012b; Golumbic et al., 2013). In seminal work, O’Sullivan et al. (2015) used a linear regression model to reconstruct speech envelopes from continuous multi-channel EEG responses to natural speech. When selectively attending to one speaker in a two-talker mixture, the speech envelopes reconstructed from the EEG responses were more correlated with the envelope of the attended speech stream compared to the ignored one. This, in turn, allowed listeners’ attention to be predicted from single-trial (1 minute) EEG data (O’Sullivan et al., 2015). Later studies have demonstrated that EEG-based attention decoding with similar models can also be achieved in more realistic acoustic scenarios (Fuglsang et al., 2017; Aroudi et al., 2019), with shorter EEG data segments (de Cheveigné et al., 2021; Ciccarelli et al., 2019; Thakkar et al., 2024), and in older listeners with hearing loss (Fuglsang et al., 2020).

These successes obtained in an offline setting with sustained attention tasks raise hope for a real-time implementation where the brain-decoded signal steers the acoustic feedback to the listener to facilitate comprehension (e.g., in a hearing aid). In a real-time system, EEG decoding is coupled with an acoustic speech separation system that separates different voices in the listening space. The decoded attention signal controls the relative gains of the separated channels, enhancing attended sources and suppressing interfering ones. The change in the relative level of the speech sources influences the neural response (Ding and Simon, 2012a; Das et al., 2018), creating a closed feedback loop (see Fig. 1). This closed loop, of which the user is a part, immediately raises several challenges. For instance, if an ignored speech stream has been attenuated by the BCI, can the user then switch attention to that stream? Is there a minimum amplitude below which this is no longer possible? Does visual feedback affect this situation, for example by helping to decode a stream when its audibility is low? How tolerant are users to inevitable processing latency and errors? How accurate and fast must the device be to be usable in real-life communication? How should the acoustic signal be rendered to the user (spatialization, etc.)? How can the acoustic speech separation strategy be integrated within the small form factor of a hearing aid? Should the BCI seek to enhance attended sources or suppress ignored ones? Does the benefit require training, and can typically elderly hearing-impaired people adapt to the device and learn to harness its power?

**Figure 1:**
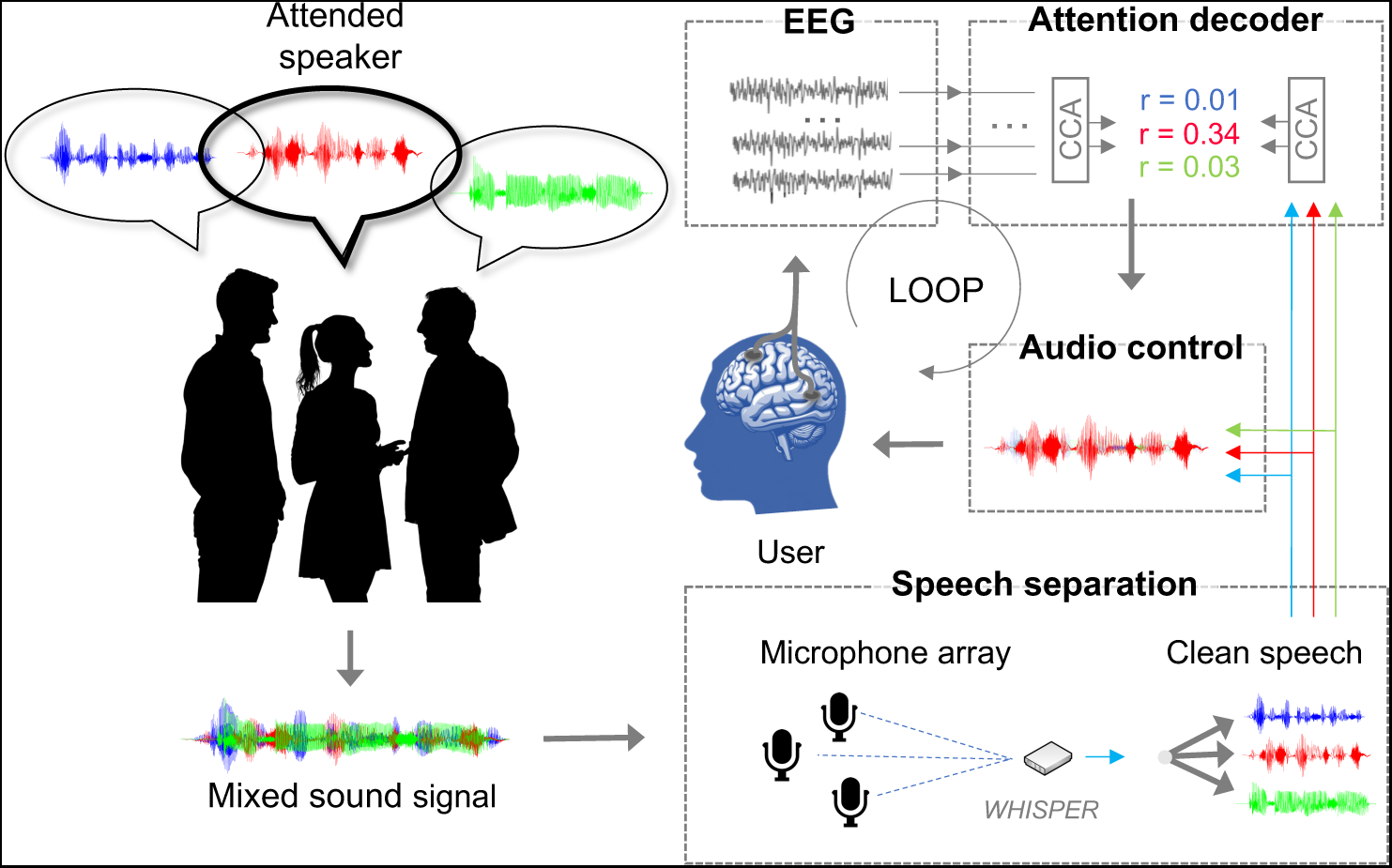
The cognitively controlled hearing aid in overview. Speech is picked up by an *ad hoc* array of microphones and processed to produce clean speech streams, one per speaker. Continuous EEG signals from the user are synchronized with the separated audio streams and transmitted to an attention-decoding module that classifies the attended speech stream based on correlations between audio and EEG features. The time-varying classification outputs are then used to control the relative gains of the separated audio streams.

Here we present a real-time implementation of such a closed-loop auditory attention decoding system. Our system addresses each requirement of a real device (hardware, software, usability) as explored within in the COCOHA project (https://cocoha.org/). It consists first of a wireless multi-microphone hardware platform that performs online low-latency acoustic speech separation via spatial beamforming. Next, an attention decoding module performs online analysis of the correlation between speech stimulus features of the separated speech streams and continuous EEG signals. By correlating the EEG to envelope features of each speech stream in a mixture, the decoder identifies the attended stream as the most correlated one. The decoded attention signal is then used to steer the relative gains of the separated speech streams to enhance the attended speaker in the acoustic feedback to the listener.

Real-time attention decoding was pioneered at the Telluride Neuromorphic Engineering Workshop in 2012, which inspired several of the offline studies mentioned above (O’Sullivan et al., 2015). More recent attempts have focused on online decoding (Zink et al., 2016; Aroudi et al., 2021) or hardware implementation (Ha et al., 2023). Our system, completed in 2018, presents an ambitious attempt to integrate hardware components for speech separation and brain decoding into one system (Wong et al., 2018b). The current paper presents an overview of this work. In section 2 we describe the various components of the system in overview. Section 4 shows demonstrations of real-time gain steering of competing audio signals in listeners with hearing impairment and with visual inputs of the speakers face. Details of the implementation and the demonstrations are given in section 3. Finally, we discuss the limitations and prospects of the cognitively controlled hearing technology in a broader context in section 5.

## 2. THE REAL-TIME SYSTEM IN OVERVIEW

### 2.1. Overview of the system

Figure 1 shows an overview of the real-time system. First, mixed sound signals in a free-field acoustic environment are picked up by a wireless microphone array placed in the listening space. A hardware platform synchronizes microphone signals between each other and performs low-latency beamforming to output separate speech streams corresponding to *N* different speakers in the listening space (section 2.2). Concurrently with this audio processing, EEG signals are recorded from the listener’s scalp and synchronized with the audio signals. Synchronized EEG and audio signals are then streamed to an attention decoding module (section 2.3). The attention decoding module estimates which stream is currently being attended to by the listener, and this knowledge is used to modulate the relative gains of the audio channels in the mixture (section 2.4). The re-rendered audio mixture with the attended speaker enhanced is then presented to the listener via an insert earpiece (e.g., a hearing aid speaker).

### 2.2. Microphone array platform and acoustic scene analysis

Modern hearing aids are equipped with multiple microphones that allow the signal-to-noise ratio (SNR) of the sound to be improved via array processing. However, the close spacing between microphones, and distance from sources (both target and interferers) severely limits this benefit. We identified this as a serious bottleneck, and thus investigated the option of an *ad hoc* array of distributed microphones, communicating wirelessly. In an *ad hoc* array, the microphones occupy arbitrary positions, and the beamformer adapts to this geometry (in contrast to preset array geometries commonly proposed in the literature). This allows for large inter-microphone distances and potential proximity with sources.

For flexibility, we designed a microphone-equipped module capable of serving both as a remote node, and as a proximal node connected to the hearing aid (which can also function in the absence of remote nodes). This custom microphone/wireless hardware platform (called WHISPER) is described in detail in Kiselev et al. 2017 and Ceolini et al. 2020b. In brief, each module supports up to four microphones (see Fig. 2), and can be combined to form a sensor network of any arbitrary number. Microphone signals are synchronized wirelessly between modules, and the array can then be used for low-latency, high-quality speech separation via beamforming computed locally on the platform. Local computing capabilities allows the computational burden of potentially complex separation algorithms to be distributed.

**Figure 2:**
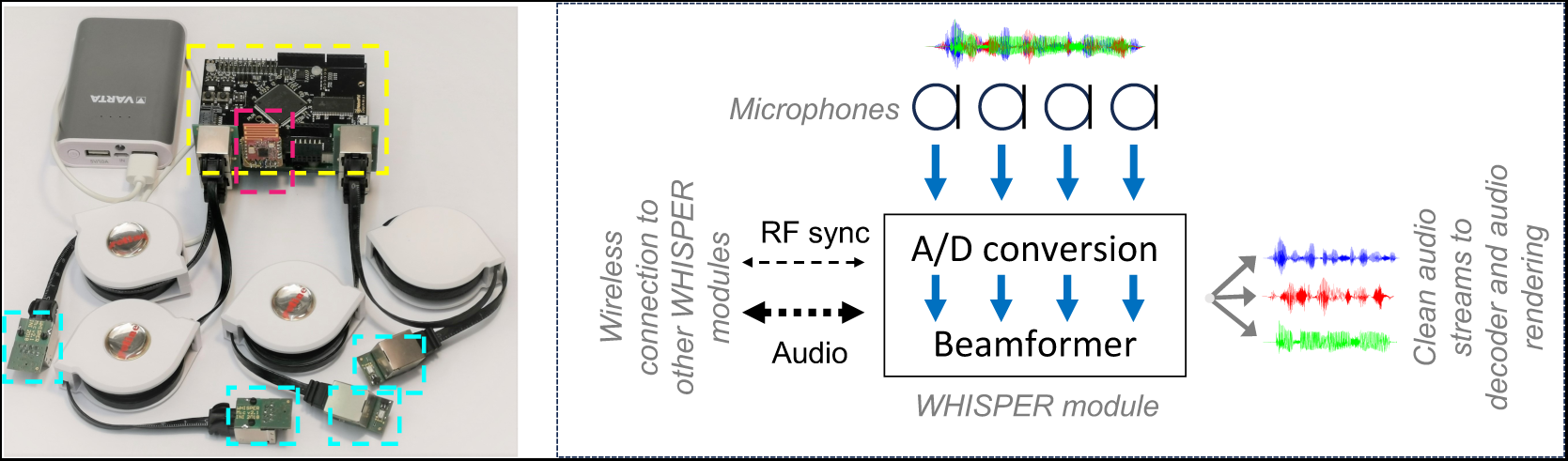
***Left*** : prototype WHISPER module (yellow box) equipped with four microphones (cyan boxes) and a wireless synchronizer (red box). ***Right*** : a WHISPER module acquires audio from up to four microphones and exchanges sampled audio and synchronization signals with other WHISPER modules or a common laptop. Microphone streams are merged and processed by a beamformer to produce clean streams for the decoder and audio rendering modules.

Theory allows for infinite interferer rejection as long as the number of microphones is at least equal to the number of sources (target and interferers). However, in the presence of noise and reverberation, or for certain degenerate configurations, a larger number of microphones may provide better performance, as observed empirically. It also offers leverage to minimize spectral distortion of the target as a result of propagation and spatial filtering.

WHISPER can be used to deploy any separation or enhancement algorithm given its ability to run floating-point algebra. For our real-time BCI, we used the simple but powerful Minimum-Variance Distortionless Response (MVDR) beamforming algorithm. In evaluations of speech separation quality within a reverberant room with 2-4 speaker mixtures (with *a priori* SNR=0), MVDR on WHISPER obtains a signal-to-distortion (SDR) of ~6 dB using 1 WHISPER node (4 microphones), and ~9 dB SDR using 3 nodes (12 microphones, see Ceolini et al. 2020b). This corresponded to a short-time objective intelligibility (STOI) score of 0.6 and 0.9, respectively. We have also evaluated WHISPER running other similar beamforming algorithms, such as the speech distortion weighted multichannel Wiener filter (SDW-MWF), maximum SNR (MSNR), masked-based MVDR (MB-MVDR) or mask-based generalized eigenvalue (MB-GEV), with equivalent high performance (Ceolini et al., 2020b). Notably, WHISPER can also be used to deploy high-performing speech enhancement algorithms based on deep neural networks (see Ceolini and Liu, 2019).

Achieving low latency is critical for successful applications in hearing technology. Two terms contribute to latency: that of the beamformer calculation (roughly 11 ms for a 512-sample buffer with 75% overlap at 24 kHz,(see Ceolini et al., 2020b)), and that of the wireless transmission protocol. WHISPER was implemented with off-the-shelf wireless protocols, the optimization of which is a target for future work. A third, negative term applies to distant sources as a result of faster wireless than acoustic transmission.

A demonstration of our BCI shown in sect. 4.1 below were conducted using a single WHISPER module (with 4 microphones) wired to a remote laptop running an MVDR beamformer. In this situation, the processing latency is essentially the algorithmic latency, roughly 11 ms.

### 2.3. Attention decoding

The real-time attention decoding pipeline is illustrated in Fig. 3. The decoder receives the separated audio streams from the microphone platform and multichannel EEG amplifier using LabStreamingLayer (Kothe et al., 2014). EEG and audio streams are synchronized via an initial trigger pulse sent along both paths before each experiment. The online processing pipeline, implemented in OpenVibe (Renard et al., 2010), then performs preprocessing of both EEG and audio streams (Fig. 3 blue), including multichannel envelope extraction for the audio signals, and standard filtering and denoising for the EEG (see sect. 3 for additional details).

**Figure 3:**
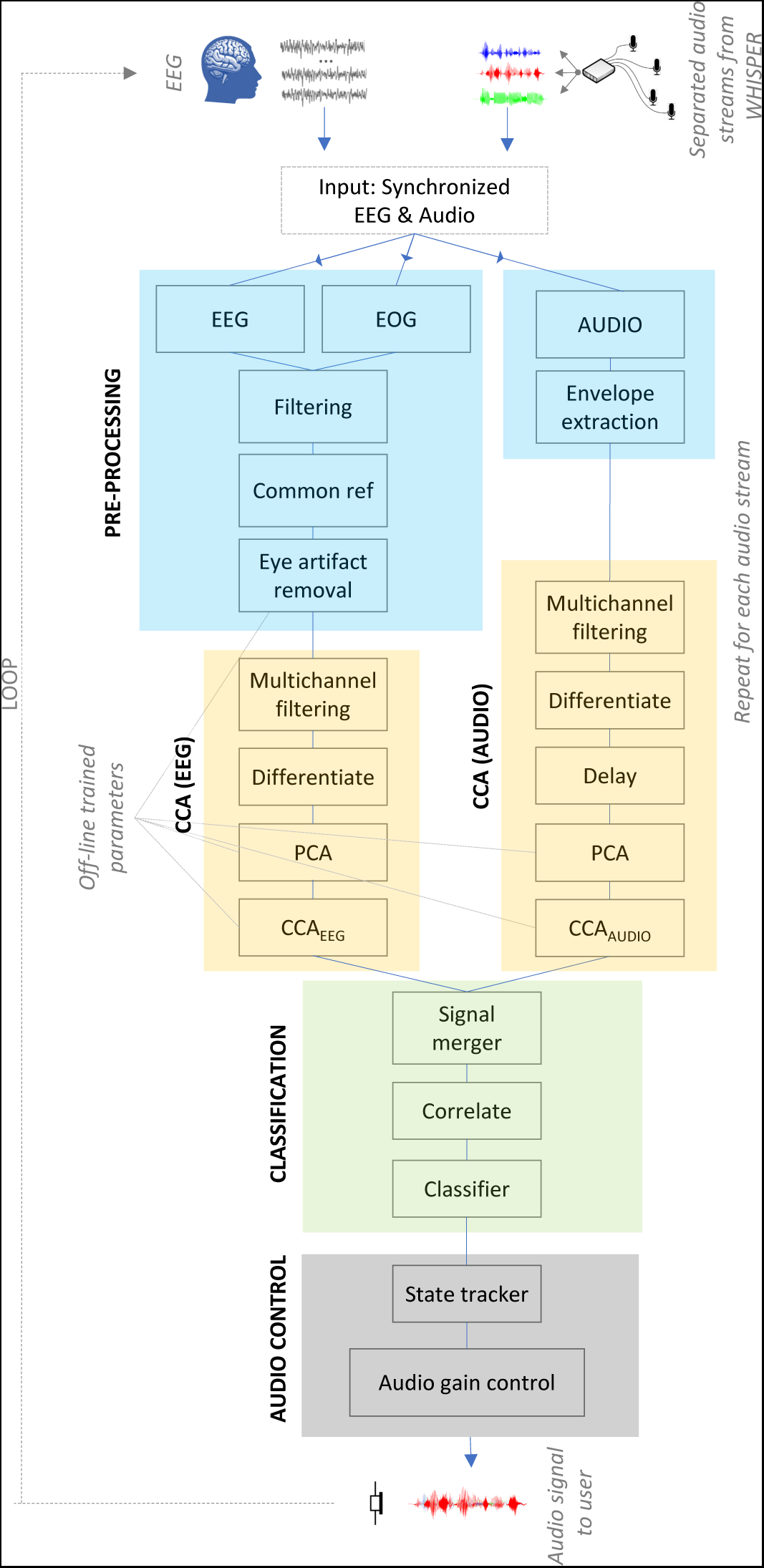
Schematic of the real-time attention decoding pipeline. Each box represents a processing object implemented in OpenVibe (Renard et al., 2010). Inputs are synchronized EEG and separated audio streams. The output is a gain control signal for each speech stream. The processing can be divided into pre-processing (blue), CCA transformation of EEG and audio (yellow), classification of attention (green), and audio control (grey).

The core of the attention classification pipeline is a real-time implementation of a linear decoder based on canonical correlation analysis (CCA) (see de Cheveigné et al. 2018 for additional details). This approach generalizes linear regression models to predict synchronized stimulus-response activity (de Cheveigné et al., 2018, 2021). The appeal of CCA over previous forward (prediction) and backward (reconstruction) regression methods is greater accuracy (de Cheveigné et al., 2018).

Based on training data, a CCA model is trained to find linear transforms that are then applied to both EEG and audio features from each audio stream at test time (Fig. 3, yellow). As inputs to the CCA model, the EEG and audio streams are passed through a multichannel filterbank that allows the CCA solution to implement filters that equalize the spectral content of both streams, and help factor out irrelevant components at multiple time scales. The filterbanks are identical on the EEG and audio side. On the EEG side, the CCA solution implements a spatiotemporal filter to capture different neural sources correlated with stimulus activity (de Cheveigné et al., 2018). On the audio side, the CCA combines envelope subbands to perform bandpass filtering that capture envelope fluctuations at different time scales (Dau et al., 1997). Once the CCA weight matrices are learned on training data, the linear transformation of the data simply consists of parallel matrix multiplications at test time (one for the EEG data, one for each separated audio channel), implying low computational cost.

The CCA-transformed EEG and audio streams are multivariate and produce sets of pairwise correlation coefficients between EEG and audio features. For each pair, the correlation scores are continuously computed within a predefined decoding window (Fig. 3, green). The duration of the decoding window represents a trade-off between decoding speed and accuracy (Wong et al., 2018a; de Cheveigné et al., 2021). In offline studies, we found that our CCA-pipeline can classify attention with an accuracy of around 70-80% with decoding windows around 6-8 seconds (see Fig. 7C below for an example of offline decoding accuracies). The windowed correlation component scores are passed to a linear classifier, here implemented as a two-class support vector machine to decide which of two streams is attended. The choice of classifier is not critical. In the simplest case, the correlation component scores can simply be summed and compared between *N* streams.

### 2.4. Audio control

The output of the attention classifier is used to control the gain of the different audio streams in the speech mixture (Fig. 3 grey). We have explored different audio control strategies. A simple first choice was to map time-varying classifier probabilities to an instantaneous audio gain on each speech stream via a non-linear mapping function. For this, we used a point non-linearity that pulls low classification probabilities towards zero, and high classification probabilities towards a maximum applied gain (see sect. 3 for details). The rationale is to leave the gain unchanged (i.e., no BCI feedback) unless the classifier has reached some level of certainty about the subject’s attention.

A drawback of a simple instantaneous coupling between classification scores and gain is the potential of erratic and fast gain changes. Even if infrequent, such acoustic artefacts may be perceived as disturbing by the user. As an alternative strategy, we implemented an attention state space model to control the audio levels (see sect. 3). In this, the rate of gain change varies with the certainty of the classifier. In each classification window, the user’s attention state is a function of the previous state and the current information transfer rate estimated from the classifier probabilities. In effect, the state tracker accumulates information over time to estimate the user’s current attention state and the relative gains change based on this state. This makes the system less prone to detrimental sudden shifts in gain upon misclassifications while maintaining the possibility of fast state changes when the current information transfer rate of the system is high.

## 3. METHODS

In this section we provide more details of the implementation and experimental methods. The busy reader may wish to skip to Section 4 for demonstrations of real-time attention steering.

### EEG acquisition

In the demonstrations shown in sect. 4 below, two different EEG acquisition systems were used. For demonstrations of the real-time platform with microphone inputs (sect. 4.1) we used mBrainTrain’s mobile 24-channel SMARTING system (mBrainTrain, Belgrade, Serbia) with a wireless DC amplifier attached to the back of the EEG cap (EasyCap, Hersching, Germany). For experiment I with hearing-impaired users (sect. 4.2) and experiment II with audiovisual stimulation (sect. 4.3), we used a 64-channel Biosemi ActiveTwo system.

### EEG training data

For all real-time experiments (sects. 4.1–4.3), offline EEG data for training the system were first obtained from each subject. Subjects were presented with two-talker mixtures (audiobooks, one male, one female) presented dichotically for 30-40 mins. The training session was split into 50 s long trials and subjects were instructed to attend to one of the two speakers in each trial. The location of the attended target (attend left vs right) was counterbalanced across trials to avoid spatial bias. In experiment II with audiovisual speech, 10 subjects received additional onscreen visual feedback of the two speakers during training, while 10 other subjects were only presented with the audio. Decoders were trained on the audio envelopes of the speech streams in both cases. In separate experiments, we noted that listening to a single speech stream during training data acquisition resulted in similar closed-loop performance. If this observation can be confirmed, this can simplify the collection of training data for attention decoding experiments considerably (de Cheveigné et al., 2021). The training data were processed offline with the same processing pipeline as described for the online system below.

### Online implementation

The real-time decoder was implemented in Python assembled as processing blocks (see Fig. 3) using the OpenVibe (1.3.0) graphical BCI software platform for real-time code integration (Renard et al., 2010). EEG and audio signals were broadcasted to an OpenViBE acquisition server via labstreaminglayer (Kothe et al., 2014), a library for networking and routing of time-series data. To synchronize the data streams, trigger pulses generated by an Arduino were sent along both paths (EEG and audio), and a receiver module in OpenViBE was then used to quantify and compensate for any delay between the streams. i.e., to maintain time-synchronization between data streams.

### Online audio processing

To extract audio envelopes in the online system, each separate audio stream was squared, low-pass filtered at 20 Hz (4^th^ order Butterworth), and down-sampled to 64 Hz via decimation. The envelopes were raised to the power of 0.3 to account for the compressive response of the inner ear (Fuglsang et al., 2020; Lopez-Poveda et al., 2003). The envelopes were further passed through a dyadic filterbank implemented as series of low-pass filters followed by a first order differentiator (de Cheveigné et al., 2018). The low-pass filtering was performed by convolving the signals with square windows of lengths ranging from 1/32 to 1 s in 10 steps on a logarithmic scale. Differentiation of the low-passed envelopes act as a high-pass filter, effectively creating a filter-bank of bandpass filters in the modulation domain. The audio envelopes were then delayed to account for the difference between stimulus and response by applying a constant shift estimated from the training data. Next, the data were transformed using principal component analysis (PCA) to reduce dimensionality. The number of PCA components retained was determined by cross-validation on the training data, and the PCA transform of the resulting dimensionality was then applied in the online processing. Finally, the multichannel envelopes were multiplied by the CCA weights obtained on the training data.

### Online EEG processing

The EEG data were low-pass filtered at 20 Hz (4^th^ order Butterworth) and down-sampled to 64 Hz to match the audio sampling rate. A high-pass filter at 0.1 Hz (2nd order Butterworth) was used to filter out low-frequency drifts, and a common average reference was applied. The data were further filtered to reduce eye-blink activity using a spatial filter obtained using the electroocular (EOG) channels in the training data (Wong et al., 2018a). The EEG were passed through a multichannel filtering operation identical to the one applied to the audio envelopes. The data were truncated using PCA, similarly as for the audio data. Finally, the data were transformed using the CCA weights obtained on the training data.

### Online audio-EEG classification

The temporally aligned CCA-transformed EEG and audio envelopes were concatenated and the correlation between corresponding CC pairs were computed within a continuously updated decoding window. Decoding windows of 8 s were used in experiment I and windows of 6.5 s were used in experiment II. In both cases, classifications where performed at a rate of 4 Hz, e.g. with a stride of 0.25 s, resulting in an decoding windows with an overlap of 7.75 and 6.25 s, respectively. Each correlation coefficient pair was z-scored based on the training data and input to a linear support vector machine (SVM) classifier with weights obtained on the training data. The classification output was converted to class probabilities for each speech stream being attended based on the training data (Platt et al., 1999).

### Audio gain control

The probability outputs of the classifier was used to control the gain balance between speech streams by attenuating ignored streams in the acoustic feedback to the listener. The baseline level of the speech mixture (without BCI feedback) was set to 60 dB HL, and BCI feedback could result in a maximum gain difference between speakers of 10 dB. For hearing-impaired listeners, the baseline level was amplified based on their audiometric thresholds to compensate to reduced audibility.

Different gain control functions were investigated. In experiment I, classification probabilities were mapped to a time-varying gain control signal *G_t_* using a non-linearity corresponding to a modified Weibull cumulative distribution function:

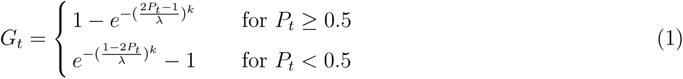

where *P_t_* is the time-varying probability outputs of the classifier. In the current two-talker scenario, speaker A is classified as attended when *P_t_ >* 0.5 and speaker B is classified as attended when *P_t_ <* 0.5 (and *P_t_* = 0.5 indicates the chance-level likelihood). The parameters *k* and *λ* determines the shape and gain of the Weibull non-linearity (see Fig. 4). The gain control signal *G_t_* was scaled to the real-valued interval [*−*1, 1] (dividing by 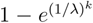). *G* = *−*1 then indicates that full gain should be applied to speaker A while speaker B should be attenuated by the maximum gain difference. *G_t_* = 1 indicates full gain applied to speaker B while speaker A is attenuated. 0 indicates that the system should not attenuate either speaker. The positive and negative values of the gain control signal were each mapped to dB and used to control the audio levels of the two speech streams.

**Figure 4:**
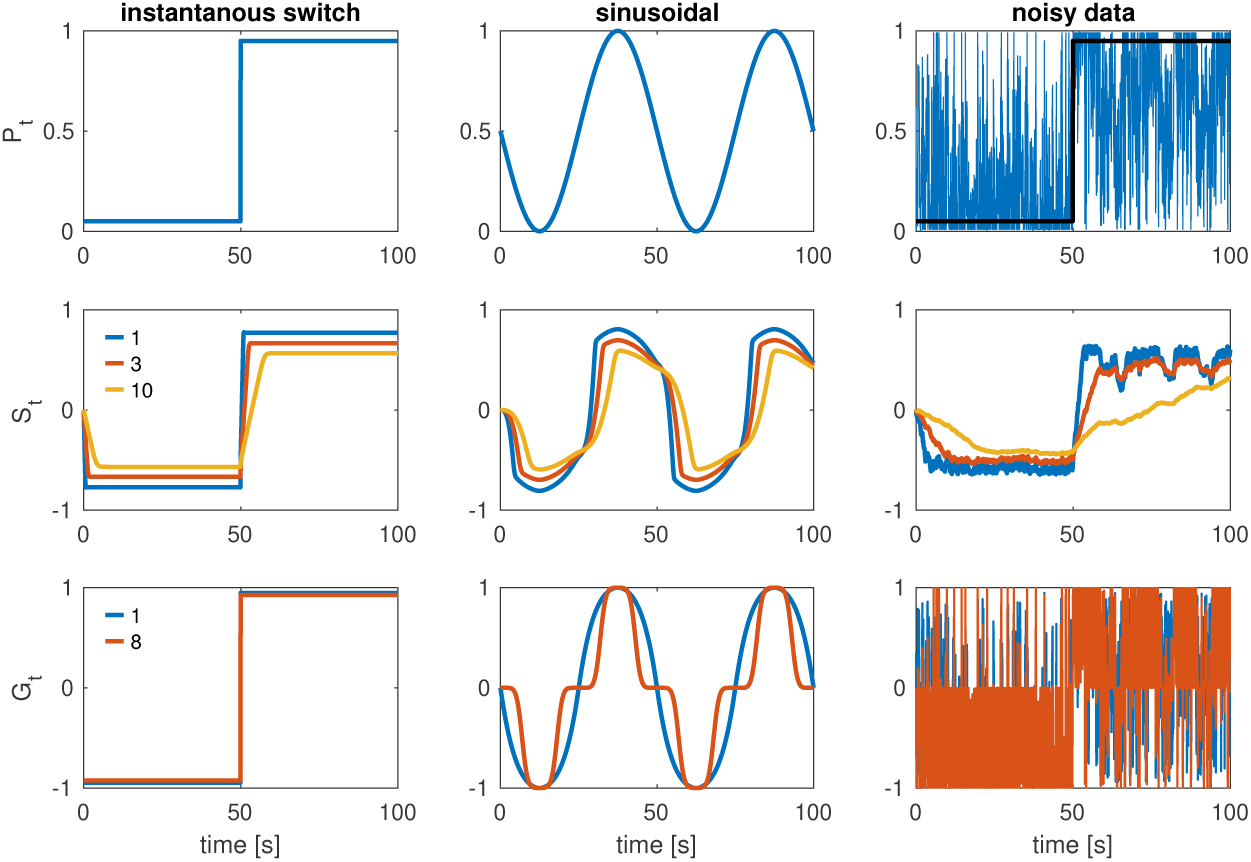
Simulations illustrating the behavior of the different audio gain control strategies. Top row: simulated attention classification probabilities *P_t_*. Middle row: outputs of the state tracker *S_t_* for different values of the parameter *K* in eq. 2, i.e., different durations of the decoding window (in secs). Bottom row: outputs of the instantaneous gain control function *G_t_* for different values of the shape parameter *k*. Three different simulations are considered here. The first simulation (left column) assumes an ideal scenario where the classifier correctly identifies an attention switch after 50 s. In the second simulation (middle column) classification probabilities fluctuates sinusoidally. The third simulation (right column) again assumes a target attention switch at 50 s but the classification is noisy. As can be seen, noisy classification results in a fluctuating gain for the instantaneous Weibull mapping function *G_t_* (bottom) but a smooth gain control for the state tracker (middle).

### State tracker

In experiment II, the relative gains were controlled via a state tracker. The state tracker implemented a strategy to control the gain based on the state of the attention decoding system. The likelihood probability of the classifier *P_t_* now controls the *rate* of change in gain. This allows the state to remain stable when the attention classifier yields a chance-level likelihood (i.e., likelihoods toward 0.5), making less prone to detrimental sudden shifts in gain.

The state at time step t is computed as:

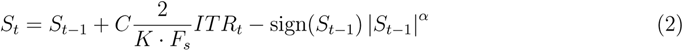

where *C* is the attention state corresponding to the classified speaker:

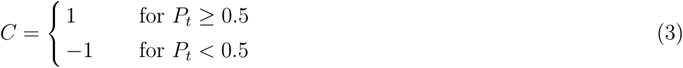

with *P_t_* again being the classifier likelihood. *ITR_t_* is the estimated information transfer rate (Wolpaw et al., 1998) in bits per classification window of length *K* seconds and *F_s_* is the number of classifications per second, here set to 4 Hz. Using ITR allows the system to adapt its switching speed to the current information rate. In essence, in absence of the sign(*S_t−_*_1_) *|S_t−_*_1_*|^α^* term, 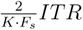 allows the state to switch from −1 to 1 (or vice versa) in one second if the information transfer rate is equivalent to 1 bit/s throughout the entire second. The information transfer rate (in bits per classification window) is computed as:

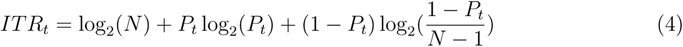

where *N* is the number of classes (2 in this case). The exponent *α* in eq. 2 acts as a hyperparameter that controls the amplitude of state changes and was fixed heuristically at 7.5 in experiment II.

Fig. 4 shows simulations that illustrate the behavior of the two different gain control strategies. The top row shows simulated classification probabilities *P_t_* in three different scenarios (columns). The resulting outputs of the state tracker *S_t_* and the instantaneous gain function *G_t_* are shown below in the middle and bottom rows, respectively. The first (leftmost) column simulates an optimally classified instantaneous switch of attention half-way through the trial. The instantaneous gain function (bottom row) shifts the gain accordingly whereas the state tracker (middle row) introduces a delay proportional to the duration of the decoding window. The middle column illustrates how each gain function maps classification probabilities, here simulated as varying sinusoidally. For higher values of *k*, the non-linear mapping in the instantaneous gain function (bottom row) will force the gain to remain unchanged for low classification probabilities and apply maximum gain difference once a certain classification level is reached. The parameter *k* controls this non-linearity and its setting allows a trade-off between avoiding gain changes for low probabilities and the overall smoothness of the gain changes. The parameter *k* mainly affects the perceptual quality of the gain track and was set heuristically (*k* = 2) in experiment I.

A main motivation for a state space approach to gain control can be illustrated with less-than-optimal classification. A noisy classification of an ideal switch (black) is simulated in the rightmost column in Fig. 4. In this case, an instantaneous gain control (bottom row) produces a fluctuating gain, whereas the state tracker (middle row) is less prone to these variations. The duration of the decoding window can be increased to make the state tracker less prone to misclassifications but at the expense of slower change in gain upon actual switches in attention. Moreover, the state space model adapts switching time based on the input ITR, i.e., the system can react faster when the current attention classification accuracy is higher. Since the noisy and variable classification scenario is closer to reality for classification based on short EEG segments, we find that a state space approach to gain control like this is generally preferable. In experiment II using the state tracker, we fixed the decoding window to 6.5 seconds to represent a trade-off between switching speed and random gain fluctuations. The duration of the decoding window could be automatically optimized for performance on training data, but this can result in very long duration windows for poor subjects, effectively making the BCI unresponsive.

### Participants

In experiment I with hearing-impaired listeners (sect. 4.2), 3 young normal-hearing and 3 older hearing-impaired listeners participated. The hearing-impaired subjects participated without their hearing aids on. Instead, a linear frequency-specific amplification was applied based on their audiometric thresholds to restore baseline audibility. In experiment II with audiovisual speech (sect. 4.3), 20 young normal-hearing listeners participated. Experiments were conducted in accordance with protocol H-16.036.391 approved by the Science Ethics Committee for the Capital region of Denmark.

### Closed-loop test procedures

In test sessions with the real-time system, 24 trials (16 closed-loop, 8 open-loop) of 90 s were collected per participant in experiment I (sect. 4.2), and 36 trials (24 closed-loop, 12 open-loop) of 50 s were collected in experiment II (sect. 4.3). Pre-recorded two-talker speech audio streams (one male, one female) were fed directly to the real-time decoder in these experiments. Subjects were prompted to attend to one speaker via on on-screen arrow at the beginning of the trial, and to switch attention to the other speaker half-way through the trial.

### Code

A software implementation of the real-time system, including code for the real-time decoding pipeline is available via https://cocoha.org/the-cocoha-matlab-toolbox/

## 4. DEMONSTRATIONS

In this section, we report different real-time experiments performed to demonstrate that the system is functional and usable by a real subject. The demonstrations serve as proof-of-principle use-cases rather than quantitative evaluations of the BCI. First, we show a video demonstration illustrating the system with the WHISPER microphone platform separating speech in a real-world acoustic scenario (sect. 4.1). Next, we show two experiments to demonstrate attention-steering across subjects with or without hearing impairment (experiment I, sect. 4.2), and with and without visual input of the speakers face (experiment II, sect. 4.3). In these experiments, rather than using the microphone platform, we used pre-recorded speech (audiobooks) so that the same speech material could be presented and results compared across different subjects. The real-time system was then used to control the audio gain of the two speech streams.

### 4.1. Demonstration with real talkers

A video illustrating the real-time system is shown below (Fig. 5). A listener wearing a wireless EEG system focuses attention on either the male or female speaker. The two speakers alternately raise their hands to indicate that the listener should attend to her or him. The WHISPER microphone platform is placed on the table between the talkers and the listener in an *ad hoc* arrangement. The microphone signals are synchronized on the platform and streamed to the speech separation module that performs online beamforming. The separated speech streams are transmitted to the attention decoding module running on a remote laptop. The decoding module receives and synchronizes EEG and audio streams and performs CCA-based decoding of attention, as described above. The classification output controls the relative gain of the two speech streams in the acoustic feedback presented to the headphones of the subject. The audio track of the left video contains the recorded room audio (the mixed speech). The audio track of the right video contains the beamformed audio presented to the subject, i.e., emulating the hearing aid output. More video demonstrations are available at cocoha.org/2018/12/31/videos/.

**Figure 5:**
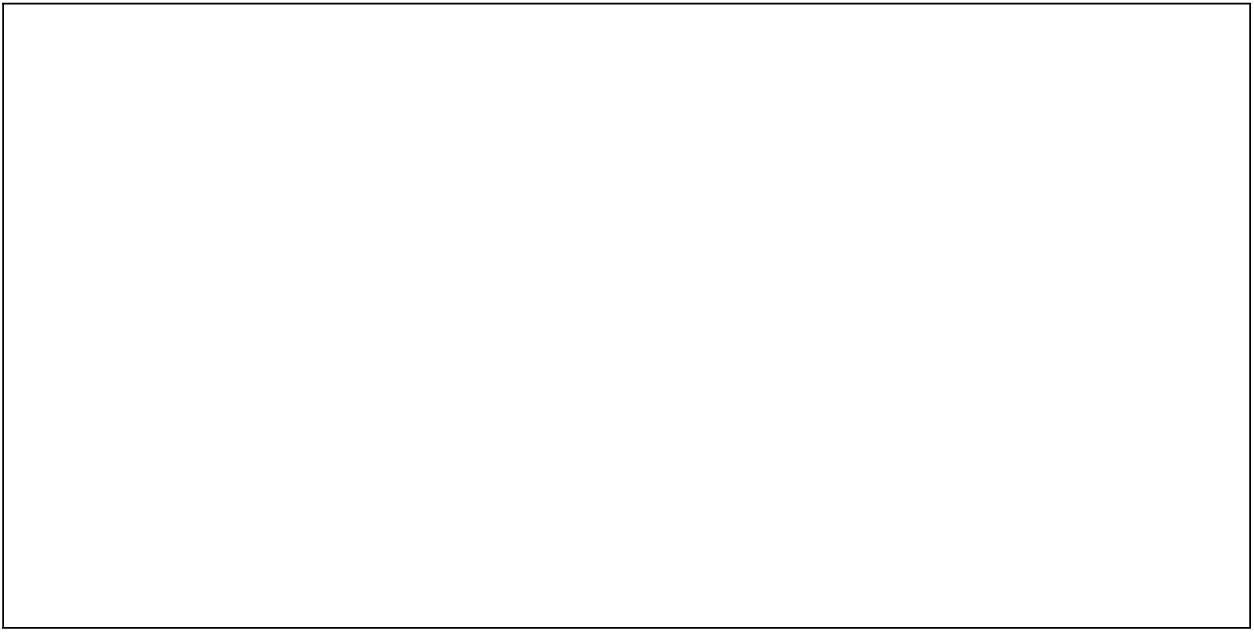
[Videos removed in this preprint. Please find the videos at cocoha.org/2018/12/31/videos/] Video demonstration of real-time attention decoding. Left: the mixed speech audio recorded in the room. Right: the separated speech audio received by the subject. The subject (facing away from the camera) is wearing a wireless EEG system and headphones receiving the acoustic feedback from the BCI. The two speakers raise their hand alternately to indicate that the listener should attend to them. The WHISPER microphone platform is placed on the table between the listener and the speakers

### 4.2. Experiment I: Demonstration with normal-hearing and hearing-impaired listeners

We first tested the real-time system with the clinical target group: older listeners with hearing impairment. Hearing-impaired (HI) listeners with moderate-to-severe hearing loss and normal-hearing (NH) listeners were presented with two competing speech streams in an attention-switching paradigm. To be able to compare results across listeners, pre-recorded speech streams (one male, one female) were presented via loudspeakers to the left and right of the subject. Training data were first obtained with subjects attending one of the two simultaneous speakers on separate 50 s long trials for ∼30 min (as in offline studies). After training the attention decoder, subjects performed closed-loop steering on 16 test trials, each 90 sec-long. 8 open-loop test trials (without neurofeedback) were collected for comparison. Subjects were prompted to attend to one speaker at the beginning of the trial and to switch attention to the other speaker after 45 s. Online classification was performed in decoding windows of 8 s at a rate of 4 Hz. The classification probabilities was then mapped to an instantaneous gain controlling the balance between the two recorded audio streams (see sect 3). Maximum classification probability resulted in a maximum gain difference of 10 dB between the two speech streams.

Data from these real-time experiments are shown in Figure 6. The average gain applied to the two speech streams as a result of real-time decoding are shown in Fig. 6C and were similar for the HI (blue) and NH (red) listeners. At the onset of trials, the two speakers are presented at the same baseline level (0 dB target-to-masker ratio, TMR). As expected from offline decoding results, the system can correctly classify the attended speaker and the acoustic gain changes accordingly at the onset of the trial. When cued to shift attention, the subject attends to the other speaker. At that point, the target speaker has a negative target-to-masker ratio, here the maximum gain difference of −10 dB TMR (supposing that the system classified the speaker correctly on the first half of the trial). As can be seen, the gain successfully switches to enhance speaker 2. The average switch time, computed as the duration from the switch cue to the zero-crossing point in the classifier output, was found to be 4.7 s (min=2.9 s, max=6.3 s).

**Figure 6:**
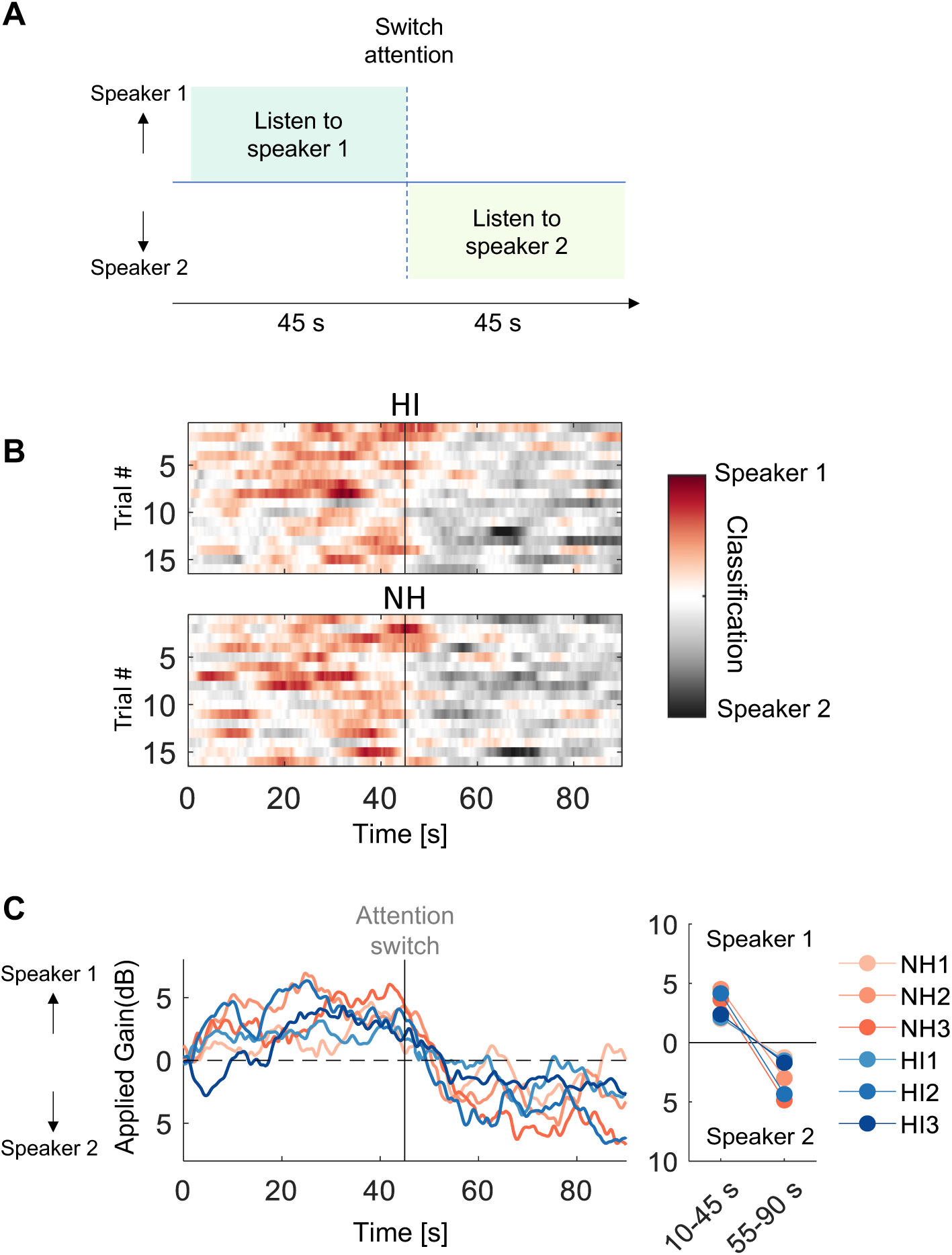
Real-time attention steering of acoustic feedback in hearing-impaired (HI) and normal-hearing (NH) listeners. (**A**) Paradigm: subjects were presented with two simultaneous speech streams and cued to switch attention to speaker 2 after listening to speaker 1 for 45 s. (**B**) Outputs of the attention classifier as a function of time. The line indicates the point of the attention switch. (**C**) Applied gain (in dB) to speaker 1 and speaker 2 as a function of time. Red traces indicate NH listeners, blue traces indicate HI listeners. The panel on the right summarizes the average gain applied across trials 0-35 s before and 10-35 s after the attention switch, i.e., omitting the 10 s following the switch.

### 4.3. Experiment II: Real-time experiment with audiovisual speech

Auditory attention can be decoded while listeners only hear the speaker they are attending, but listeners typically look at the speaker they are listening to in real-world situations. It is unclear whether a BCI trained only on auditory inputs might also work successfully in natural audiovisual conditions. The system could potentially fail to generalize to this mismatching stimulus condition. Alternatively, the visual face may be correlated with audio envelope information (O’Sullivan et al., 2017a; Pedersen et al., 2022), and it may be easier to focus auditory attention on a speaker that can be seen, which may in turn improve decoding (O’Sullivan et al., 2013). Potential visual benefits are important to investigate, for instance in the case were the user wants to switch attention to a previously ignored speaker. In this case, visual information correlated with the speech signal may help ‘recover’ a target with low audibility.

We tested these questions in a larger experiment. Training data were obtained from normal-hearing participants (*N* = 20) listening to two-talker speech mixtures with or without watching the visual input of the speakers’ moving faces (Fig. 7). The stimuli consisted of two separate pre-recorded speech videos, i.e. the microphone platform was not used in this experiment. During training trials, half of the participants (*N* = 10) received audiovisual stimuli, while the other half (*N* = 10) only received the speech audio. All listeners were then tested using the closed-loop system either with audiovisual stimuli or with the speech audio alone. As a control condition, we included trials without BCI control, i.e. trials where the audio gains were left unchanged, referred to as ’open-loop’ trials. All participants were presented with audiovisual stimuli in the open-loop trials. In test trials of 50 seconds, subjects were prompted to follow one speaker and then cued to switch attention midway through the trial (i.e., after 25 s), as in experiment I. Rather than the instantaneous gain control function used in experiment I, the audio gains were controlled by a state tracker with a sliding decoding window of 6.5 s and a classification rate of 4 Hz. The rationale of the state tracker is to make the gain less prone to erratic changes in gain with noisy classification outputs (see sect. 3 for a comparison between gain control strategies). The same speech envelope decoding pipeline as described above was used in both audiovisual and audio-only conditions (i.e. no visual features were used in the decoder).

**Figure 7:**
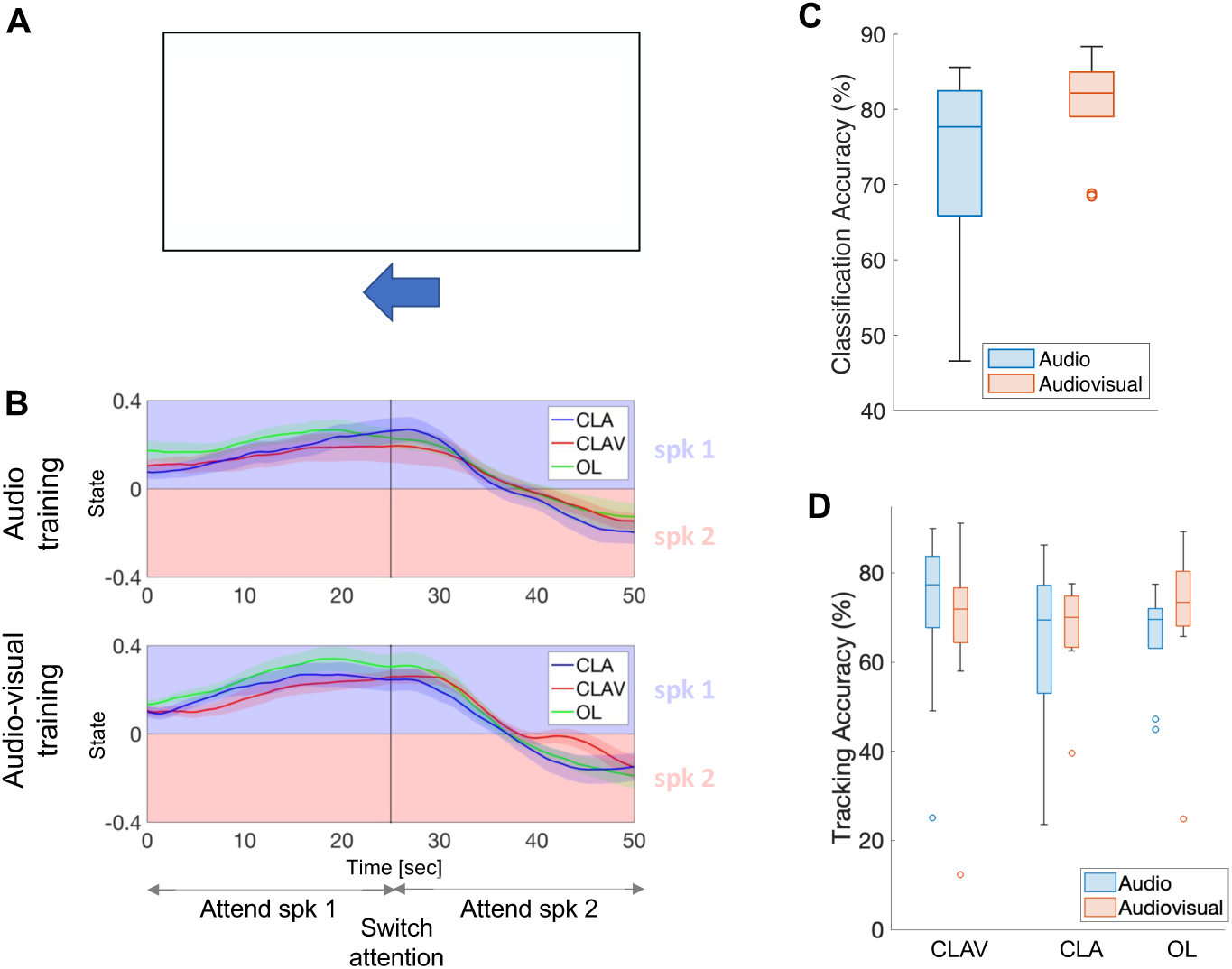
Experiment II: real-time attention decoding with and without visual input. (**A**) Listeners were cued to attend a male or female speaker in audiovisual or audio-only conditions. [Photographs removed in preprint]. (**B**) Real-time outputs of the attention state tracker for listeners trained with audio-only stimuli (above) or listeners trained with audiovisual stimuli (below). Line colors indicate the different BCI stimulus conditions. Blue: closed-loop audio-only (CLA), red: closed-loop audiovisual (CLAV), green: open-loop. The shaded areas indicate standard errors of the mean. (**C**) Offline average attention decoding accuracies based on analysis of the training data in segments of 6.5 s for listeners trained with audio-only stimuli (blue) or listeners trained with audiovisual stimuli (red). (**D**) Online accuracies of the attention state tracker for listeners trained with audio-only stimuli (blue) or listeners trained with audiovisual stimuli (red).

Figure 7 shows data from the real-time experiments with audiovisual speech. Fig. 7B shows the output of the attention state tracker in the different BCI conditions (closed-loop audio-only, closed-loop audiovisual, open-loop) for the participants trained with audio-only speech (above) and the participants trained with audiovisual speech stimuli (below). Participants were similarly able to steer the BCI with and without seeing the speakers face (CLA vs CLAV), and with and without having seen the speakers face during training. The average switching time of state tracker, again computed as the duration from the switch cue to the zero-crossing point, was found to be 12.6 s, with no difference found between BCI conditions. This slower switching time compared to the ones observed experiment I (sect. 4.2) indicates that more stable gain variations with the state tracker comes at the expense of a slower reactivity of the BCI. Fig. 7C shows the attention decoding accuracy obtained with the offline training for participants listening to audio-only speech (blue) and for participants presented with audiovisual speech (red). A modest improvement with visual feedback was seen in the offline training data, but this did not appear to translate to the real-time performance. As a summary measure of BCI tracking performance, Fig. 7D shows the average accuracy of the differential output of the state tracker, i.e., whether the state tracker steps in direction of the target speaker in each classification window. No differences between closed-loop vs open-loop steering were observed, indicating that subjects can switch attention and ‘recover’ an attenuated (previously ignored) speaker. We also did not observe interactions between training modality (audio-only vs audiovisual) and closed-loop modality. It might be expected that participants presented with audiovisual speech during training (red) performed better in the matched audiovisual closed-loop condition, and vice versa for the audio-only group (blue), but this was not observed. While the real-system accurately tracks attention for most subjects, outlier subjects performing at chance-level can be observed (Fig. 7D).

## 5. DISCUSSION

Helping hearing-impaired listeners follow speech in noisy everyday environments remains a fundamental challenge for hearing technologies. In this paper, we provide an overview of a real-time system for acoustic speech separation controlled by attention. As proof-of-principle, we describe initial closed-loop demonstrations indicating that listeners, including elderly people with hearing loss, can steer a gain to enhance the speech sources they are focusing their attention on. If the SNR of speech sources can be directly controlled and improved in this way, a gain in speech understanding follows inherently. Yet, these results should be interpreted with caution. Numerous challenges in bringing such technology to real-life use obviously remain. In the following, we discuss some of the critical prospects and limitations.

### Using attention for seamless BCI control

Using attention to steer acoustic feedback has inherent benefits in the context of neural prosthetics. BCIs often require the user to direct attention to the BCI device itself (e.g., as in a ”P3-speller” where the subject actively focuses on onscreen letters to spell words). The attention-controlled hearing instrument, on the other hand, decodes and ‘amplifies’ a natural listening process. Listeners naturally orient attention towards relevant sources in real-world listening situations, allowing them to better perceive relevant sources and suppress background noise. Our BCI assists this natural behavior without interrupting it. If the system correctly decodes attention, then the interaction is seamless. An alternative solution might be to have the user control the speech separation system via an overt device (e.g., selecting which talkers to enhance on a mobile phone). However, such a solution has the detrimental effect that it diverts the user’s attention towards steering the device rather than facilitating it. With the proposed BCI, listeners simply attend to speech as they do in everyday situations to better perceive the relevant source.

### BCI based on stimulus-response correlation

A real-time auditory attention decoding system based on stimulus-response correlation is unique in a BCI context. Most existing BCI systems classify brain signals to generate some form of system command (e.g., steering some application), i.e., the neural data are the only input to the BCI. The BCI discussed here, on the other hand, takes both EEG and audio signals as inputs. Classification now relies on attention-related changes in the *relation* between speech stimulus features and the EEG response. This imposes an important constraint on the classification problem: the BCI can act specifically on the part of the neural response that is related to the stimulus and not on global changes in brain states. This may make the BCI less susceptible to random EEG noise or task-unrelated neural signals.

An alternative solution is to classify a listener’s spatial attention based on changes in EEG alpha power (Kelly et al., 2005; Geirnaert et al., 2020; Popov et al., 2023), i.e., classifying spatial attention based only on EEG features. Yet, this solution suffers from the inherent problem that changes in lateralised EEG alpha power coincide with many task-unrelated changes in the cognitive state of the subject. In pilot studies using this approach, we found that high attention classification accuracies (*>* 90%) could be achieved with short EEG segments (*<*1 s), but the system failed to work robustly in closed-loop. A BCI based on stimulus-response metrics may help the critical generalization to real-world conditions beyond those used for training the system.

Other work has investigated neurofeedback based on auditory attention decoding with stimulus-response models. Zink et al. (2016) provided subjects with visual feedback of online decoding accuracy scores. In a system similar to ours, Aroudi et al. (2021) used stimulus-response-based attention decoding to steer the relative gain of two speech audio signals. Consistent with our observations reported above, closed-loop decoding accuracy was found to be on par with open-loop decoding (without neurofeedback). This is encouraging, given that closed-loop gain reduction of the ignored speaker could make attention switching more difficult.

Our experiments indicate that it is possible to ‘recover’ previously ignored sources attenuated by up to 10*dB* relative to the previous target source, but the boundary conditions for this remain to be explored (Das et al., 2018). If the stimulus-response relation is not detectable (at very low SNRs), then the BCI cannot classify attention. This may represent a potential weakness of decoding with stimulus-response models. However, offline experiments with audiovisual speech tracking indicate that ‘envelope’ speech tracking may be possible in silent lipreading conditions where the speaker can only be seen (O’Sullivan et al., 2017a). Visual feedback of the speaker may thus help a BCI track a speech source even when it cannot be heard.

### What is accurate decoding?

Attention classification implies a trade-off between speed and accuracy. A critical limitation of the technology is a very low tolerance for misclassification. In our experiments with hearing-impaired listeners (sect. 4.2), subjective reports from participants indicated that misclassifications were experienced as highly disturbing. With decoding windows below 10 seconds, the accuracy is currently below 100% correct classification for most subjects. This implies that the system occasionally amplifies the unattended source and suppresses the attended target, which is the opposite of the intended effect. Even if misclassification occurs infrequently (e.g., 10% of time time), the effect may be disastrous. While the BCI is seamless when classification is correct (the subject simply listens to speech), incorrect classification immediately makes the listener aware of the presence of the BCI. In our experiments, some test participants reported that this meant switching strategy to actively steering the device, a process reported as effortful, i.e., the opposite effect of the intended purpose of the BCI. However, in our real-time experiments, such subjective reports were not collected in a systematic manner. Interestingly, Aroudi et al. (2021) reported subjective ratings of perceived effort but found no effect of closed-loop steering. Yet, it is not *a priori* clear how decoding accuracy should translate to the subjective perception of BCI control or perceived effort, and examining subjective reports systematically will be important for future closed-loop studies.

As noted in the reported experiments, between-subject variability in real-time decoding accuracy is considerable. Although average performance was relatively high, some participants showed real-time tracking accuracy well below chance level (Fig. 7D). This implies that the BCI effectively works against these subjects (amplifying the speech stream to-be-ignored). The reasons for this behavior are difficult to elucidate. It is possible that some participants become aware of the feedback control and focus on suppressing the speaker to be ignored. It is also possible that occasional amplification of the ignored speaker may cause to the ignored speech to grab the subject’s attention, effectively causing the BCI to get stuck in the wrong state. Identifying such sources of failure in individual users will be critical to the success of such closed-loop technologies. In the reported experiments, we focus on showing the gain change after the instructed attention switch, but this is arguably a limited metric to capture BCI system performance. A multitude of system design choices can influence BCI performance in ways that are difficult to predict. For instance, different decoding methods or gain control strategies define the ‘reactivity’ of the BCI, which may in turn influence subjects’ attention, influencing decoding performance, etc., in a complex loop of events. Non-linear neuro-behavioral feedback systems like this requires dedicated tools of analysis and experimental setups that is beyond the scope of this study.

It is unclear how closed-loop attention steering can best be evaluated in more natural situations while preserving proper experimental control. For simplicity, and for experimental control, we limit ourselves to simple two-talker setup with access to clean audio sources. The same two-talker setup is used to generate data for model training and for closed-loop testing, and it is not clear how performance generalizes beyond this artificial situation, e.g., to different acoustic conditions, different SNRs, different number of speakers, etc.

### Better attention decoding?

Improving decoding accuracy may be criticial for progress towards a successful technology. Our current implementation relies on linear mappings (CCA) between audio and EEG features. Non-linear extensions have been proposed in offline attention decoding studies (de Taillez et al., 2020). An obvious extension is to consider deep learning-based models (Ciccarelli et al., 2019; Vandecappelle et al., 2021; Thornton et al., 2022). However, a number of pitfalls, often deceitfully hidden, should be pointed out. One concern is the limited availability of large amounts of unbiased training data. In typical attention decoding experiments, listeners are instructed to attend to one speaker in one ear and ignore a different speaker in the other ear. Linear methods based on envelope-EEG correlation can generalize to new speakers because they explicitly model this correlation. However, a non-linear neural network trained to classify attention may simply ignore the EEG and classify acoustic differences between the speakers. To avoid this, experiments that generate training data should be carefully counterbalanced, e.g., attending equally to both speakers, equally attending left/right, etc. Even here, subtle differences may leak into the training data (Rosenblatt et al., 2024). For instance, speakers may slightly shift voice pitch when speaking over longer durations, and conventional cross-validation procedures (such as leave-one-trial-out) are prone to bias from such details.

### What is ground-truth for attention decoding?

Another critical challenge for obtaining high decoding accuracy is the lack of ground truth measures of attention. Decoding models, such as the ones presented here, are trained on data obtained with listeners cued to attend to a given target during training trials. The subjects’ attention state is assumed to remain stationary throughout a trial, but the attention state cannot be monitored in any clear way. In everyday tasks, it is difficult to maintain focused attention over time, and attention fluctuates with the internal state of the listener. Attention can also fluctuate with external salient events that occasionally grab attention (Esterman et al., 2013). When instructed to focus on a speech target in attention decoding experiments, a listener’s attention may still occasionally switch to the non-target, or attention may simply drift from the task altogether (Smallwood and Schooler, 2015; Weissman et al., 2006). The lack of measures of such attention dynamics puts decoding models on uncertain ground. An alternative approach may be to dispose of attention labels and focus instead on modelling the stimulus-response relation with simpler stimuli, e.g., listening to a single speaker. For instance, de Cheveigné et al. (2021) proposed training stimulus-response models based on whether or not a segment of EEG matches the auditory stimulus that evoked it. This approach allows models to be trained in a self-supervised manner. If attention modulates the stimulus-response mapping, then robust stimulus-response models may be key to progress without relying on attention labels.

### Need for speed?

Improved decoding accuracies can be obtained by integrating the classification over longer durations, i.e., by sacrificing classification speed. In our experiments, we explored different gain control strategies. Instantaneous gain (used in experiment I) allowed faster average gain changes with attention switches (*<* 10 seconds), at the expense of erratic gain changes. Temporal integration with a state space model (in experiment II) stabilized gain fluctuations, but at the expense of slower mean switching times (*>* 10 secs). Similar long switch delays were reported by Aroudi et al. (2021). In dynamic communication situations involving many talkers, attention may switch rapidly and decoding windows of several seconds are already unfeasible. However, in other situations, longer time constants may be acceptable. In many communication situations, the communication partners remain the same over longer periods – e.g., in a restaurant or at a meeting - where the BCI can slowly ‘zoom in’ on relevant sources. An alternative decoding strategy may be to identify such long-term stable communication contexts.

### Alternative EEG solutions

A critical limitation of the technology is the quality of EEG signals recorded during natural behavior. Cortical signals recorded with scalp EEG are inherently weak and noisy. Our current demonstrations of a real-time BCI are still limited to stationary setups where the listener is sitting still and passively listening. Motion artifacts during natural behavior tend to dramatically surpass the brain signals of interest in scalp EEG. Ear-EEG (EEG recorded in the ear canal or behind the ear) offers feasible solutions for integration with a hearing aid (Looney et al., 2012; Bleichner and Debener, 2017), but does not appear to improve decoding performance (Fiedler et al., 2017). Invasive cortical recordings improve brain signal SNR considerably, and O’Sullivan et al. (2017b) demonstrated fast and accurate decoding using electrocorticography (ECoG) recordings in neurological patients (O’Sullivan et al., 2017b). New multichannel implant solutions are emerging for BCI perspectives (Musk et al., 2019; Steinmetz et al., 2021). Less invasive solutions such as subcutaneous EEG are currently in clinical development (Viana et al., 2021) and offer interesting perspectives for decoding attention. However, weighing the potential benefits of invasive solutions against the risks for the individual patient is naturally a complex ethical discussion.

### Acoustic speech separation solutions

Using a separate speech separation front-end based on distributed microphones was a design choice in our system. Distributed sensors on dedicated hardware offer high-quality speech separation at very low latencies, which is paramount for hearing instrument applications. One drawback of stationary microphones is the need for calibration for a given spatial configuration of sound sources. However, Voice Activity Detectors (VADs) could be used for dynamic calibration in non-stationary environments. More flexible extensions have recently been proposed, such as ‘acoustic swarms’ of self-distributing microphones that dynamically locate and separate speech sources in changing environments (Itani et al., 2023). Other challenges arise with synchronization of wireless nodes based on radio transmission, which is generally poorly suited for outdoor environments or rooms with plentiful reflective materials like metal surfaces.

An obvious drawback is the need for distributed microphones in the listening space. However, solutions with user-worn microphones (placed on the hearing aid) are also conceivable. Single-channel or two-channel solutions may exploit advances in speech separation within deep learning (Luo and Mesgarani, 2018). O’Sullivan et al. (2017b) already demonstrated that attention can be reliably decoded from the outputs of a single-channel speech separation system, i.e., without assuming access to the clean speech sources (O’Sullivan et al., 2017b). Successful attention decoding does not follow automatically from previous results with ‘clean sources’ since some degree of cross-talk between the separated streams must be expected even with high-performing speech separation systems (Subakan et al., 2021; Ravanelli et al., 2021).

In our system, the speech separation front-end is separate from the attention decoding process. The task of the decoder is then to decide which of *N* separated streams the listener is attending. An alternative framework is to combine the speech separation and decoding steps into one process (Ceolini et al., 2020a). Since a stimulus-response decoder estimates speech features (typically amplitude envelopes) of the attended stream, these can be used as a ‘hint’ of the clean audio source. Ceolini et al. (2020a) presented a neural network-based speech separation system trained on the neural data to extract the clean audio signal directly from the speech mixture. Using ECOG or EEG-reconstructed speech envelopes, the system directly separates the target from the mixture input. Higher envelope reconstruction accuracies were shown to translate into higher separation quality (e.g., an EEG-based reconstruction accuracy of 0.35 corresponds to around 4.2 dB SDR as reported in Ceolini et al. 2020a). This integrated approach could potentially be expanded to end-to-end neural networks that perform speech separation directly from the raw audio mixture and the continuous EEG (Hosseini et al., 2021). An advantage of such an approach is that the speech separation system does not have to separate all sources in an acoustic scene (and compare the brain response to each acoustic source) but can focus on separating only the relevant target source from the mixture. This may offer a considerable computational advantage in complex acoustic scenarios with many sources.

### Audio control strategy

In our closed-loop experiments, we use the BCI to apply audio gain to the attended and ignored speakers. Successful development of the technology is likely to require development of refined techniques to render the acoustic feedback in ways that are intuitive and comfortable for the user. Re-rendering the acoustic scene with spatially distributed sources may be critical for real-life usability. It should also be noted that amplification is not the only possible audio control strategy. Hearing aids can, for instance, benefit greatly from selectively applying different dynamic-range compression to the target and background signals (Overby et al., 2023). Selectively applying fast-acting compression to the attended foreground signal and slow compression to the ignored background avoids the problem of amplifying noise components along with the target speech, a recurrent issue faced by hearing aids today (May et al., 2020).

### Attention training

The BCI is presented here as an assistive technology for the hearing impaired, but other uses can be envisioned. Closed-loop neurofeedback training of attention has been considered as a tool to treat attention deficit disorders, helping patients sustain focus over longer periods (Monastra et al., 2006; Arns et al., 2009; Belo et al., 2021). Closed-loop imaging designs have been proposed in which people monitor their own attention via specially designed visual stimuli that change with attention-driven neurofeedback (Debettencourt et al., 2015). Our closed-loop system decodes attention during natural listening and modifies the attended speech stimulus itself. This has the advantage of engaging a natural attention process where the ‘reward’ of successfully attending is inherent in natural behavior (listening becomes easier when attending). Task difficulty can easily be titrated (e.g., via the amount of gain applied) making the system usable in attention training programs. Closed-loop training may also be relevant in context of the proposed hearing technology. It is possible that hearing impaired users may learn to adapt to the closed-loop gain control system, and individual training programs that gradually adjust SNR to improve decoding accuracy over time could be explored.

### Looking forward

Going back to the motivations laid out in the Introduction, it is clear that we are now closer to our goal, and yet we remain far from a usable product: the glass is half full, and yet half empty. We successfully assembled and tested a real-time device that covers the full chain from sound waves and brain signals to attention-based selective stimulation at the ear. However, the demonstration remains sketchy at many points. We advanced the state of the art of auditory attention decoding, but latency and accuracy remain below levels required for comfortable use. Progress here may come from better algorithms, better brain signal cues, better understanding of the intentional processes that guide selective auditory attention, or novel brain signal measurement techniques. We designed and assembled a modular distributed microphone array solution to address the problem of delivering high-quality sound to the user. However, this is just one of a wide range of possible solutions, and latency of wireless communication remained a problem. Further progress in this part of the design space depends crucially on low-latency protocols for wireless communication, and much work remains to evaluate this approach relative to others. Our experiments with human subjects, including typical potential users, indicate that the concept is workable but limited by current performance levels. These experiments also point to the need of a more complete methodology for future studies. Overall, these outcomes demonstrate that the goal is worth pursuing, and the software that we provide may help future endeavors in that direction. This work should be seen as an early milestone along the road towards a cognitively controlled hearing aid.

## Acknowledgments

This work was supported by the EU H2020-ICT grant 644732 (COCOHA), and grants ANR-10-LABX-0087 IEC and ANR-10-IDEX-0001-02 PSL. It draws on work performed at the 2016 Telluride Neuromorphic Engineering workshop, and benefitted from the boundless enthusiasm of Shihab Shamma, Nima Mesgarani, Daniel Pressnitzer, Tobi Delbruck, and others. Alain de Cheveigné is funded in part by the FrontCog grant ANR-17-EURE-0017. Søren A Fuglsang is funded in part by the grant NNF17OC0027872 from the Novo Nordisk Foundation. Søren A Fuglsang, Jonatan Märcher-Rørsted, and Jens Hjortkjær are funded in part by the Center for Auditory Neuroscience supported by the William Demant Foundation.

